# Single-shot light-field microscopy captures delivery-dependent dye and autofluorescence patterns in *Caenorhabditis elegans*

**DOI:** 10.64898/2026.09.07.749827

**Authors:** Robert Markus, Edward Rea, Nana A. Berfi, Vincenzo Taresco, Chris Hudson, Dominic R.A. White, Veeren M. Chauhan

## Abstract

Three-dimensional imaging of live *Caenorhabditis elegans* commonly relies on sequential z-stack acquisition, which can be slow and susceptible to movement and photobleaching. Here, we used light-field microscopy to capture dye and delivery-dependent fluorescence patterns across intact adult worms in a single exposure per channel. With a 40×/1.20 NA water-immersion objective, each reconstructed dataset contained 121 axial slices spanning 109 µm at 0.91 µm spacing. In a representative adult, nematode signal extended from 40.88 to 85.4 µm, corresponding to approximately 44.5 µm of captured axial depth without sequential z scanning. Three fluorescence channels were acquired with a summed exposure time of 112 ms across a 685 × 1010 µm field of view. FM1-43 labelled intestinal, cuticular and vesicular structures, FM4-64 highlighted ingested bacteria and intestinal compartments, and Nile Red revealed vesicular and broader whole-organism fluorescence. Intrinsic blue and green autofluorescence signals were also resolved. These observations demonstrate rapid single-shot acquisition of whole-organism three-dimensional fluorescence information and show how delivery route influences the spatial distribution of commonly used fluorescent probes in *C. elegans*.

**Highlights:**

- Light-field microscopy captures whole-worm 3D fluorescence in a single exposure
- A 109 µm axial range is rendered as 121 slices at 0.91 µm spacing
- FM1-43, FM4-64 and Nile Red show distinct volumetric staining patterns
- Delivery route alters fluorescence distribution across intact *C. elegans*
- Three-channel volumetric imaging uses a summed exposure time of 112 ms

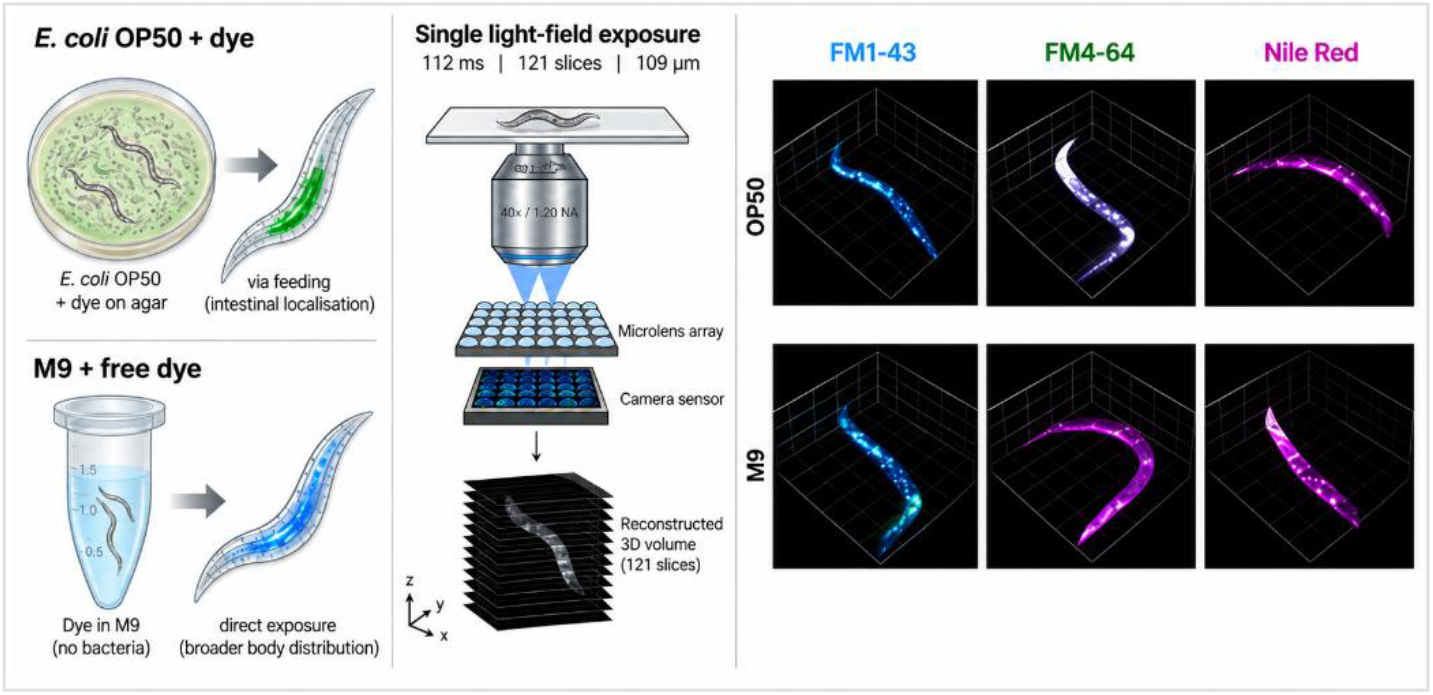

## Introduction

Volumetric imaging of live Caenorhabditis elegans is typically conducted through sequential optical sectioning followed by reconstruction of a z-stack. Although effective, this workflow is relatively slow and is vulnerable to motion artefacts, photobleaching and practical limits on throughput. A method capable of capturing biologically meaningful three-dimensional information from a single exposure would therefore be useful for rapid assessment of live nematodes, particularly when multiple fluorescent signals or delivery conditions are being compared.

## Results and Discussion

Light-field microscopy addresses these constraints by reconstructing volumetric information from a single camera exposure, thereby reducing acquisition time, motion distortion and overall light dose [1–3]. In the ZEISS Lightfield 4D implementation used here, the optical module simultaneously records multiple views containing spatial, angular and optically shifted focal information. Axial information is therefore encoded during acquisition and reconstructed afterwards, without mechanically stepping the objective through sequential z positions. To assess this capability in a simple live-imaging workflow, three commonly used dyes, FM1-43 [4], FM4-64 [10] and Nile Red [5], were delivered to adult wild-type nematodes by two methods. For the first method dye was soaked into an E. coli OP50 lawn and nematodes were allowed to feed on the labelled bacteria [6]. In the second method, nematodes were fully suspended in dye solution in M9 buffer. An identical treatment structure was used for all dyes within each delivery route, enabling direct comparison of staining behaviour across conditions. Due to the high sensitivity of the camera, endogenous blue fluorescence was also readily detected, consistent with previously described death-associated autofluorescence in *C. elegans* [7].

Complete volumetric datasets were reconstructed from single light-field exposures of 12–50 ms per channel (112 ms across all three channels), followed by offline reconstruction (∼59 s per dataset). With the 40×/1.20 NA water-immersion objective, each rendered volume contained 121 axial slices spanning 109 µm at 0.91 µm spacing. In a representative adult nematode, fluorescence signal extended from 40.88 to 85.4 µm of the rendered volume, corresponding to approximately 44.5 µm of axial worm depth encoded within each single-channel exposure. The z information is reconstructed from the simultaneously acquired light-field views rather than collected by mechanically stepping the focal plane through the specimen. By comparison, an equivalent tiled confocal z-stack covering the same field would take of the order of 18 min per worm (see Methods), approximately four orders of magnitude longer than the summed exposure time of 112 ms across three channels.

Collectively, these observations show that light-field microscopy captures biologically interpretable, dye-dependent volumetric patterns in live *C. elegans* using simple sample preparation. The differences between FM1-43, FM4-64 and Nile Red, as well as the differences between feeding and direct exposure, were retained in the reconstructed volumes. This indicates that the methods are well suited to rapid exploratory imaging where staining route, signal localisation and whole-worm context are all important. In addition to dye-derived fluorescence, endogenous autofluorescence was monitored. Blue autofluorescence was used to visualise signal associated with non-viable or compromised animals [7,8], while green autofluorescence was recorded as a distinct endogenous signal. These endogenous signals were resolved alongside dye fluorescence in the reconstructed volumetric datasets, extending the information obtained from a single acquisition. Representative reconstructed volumes and dye-specific patterns are shown in Figure 1.

**Figure 1.**
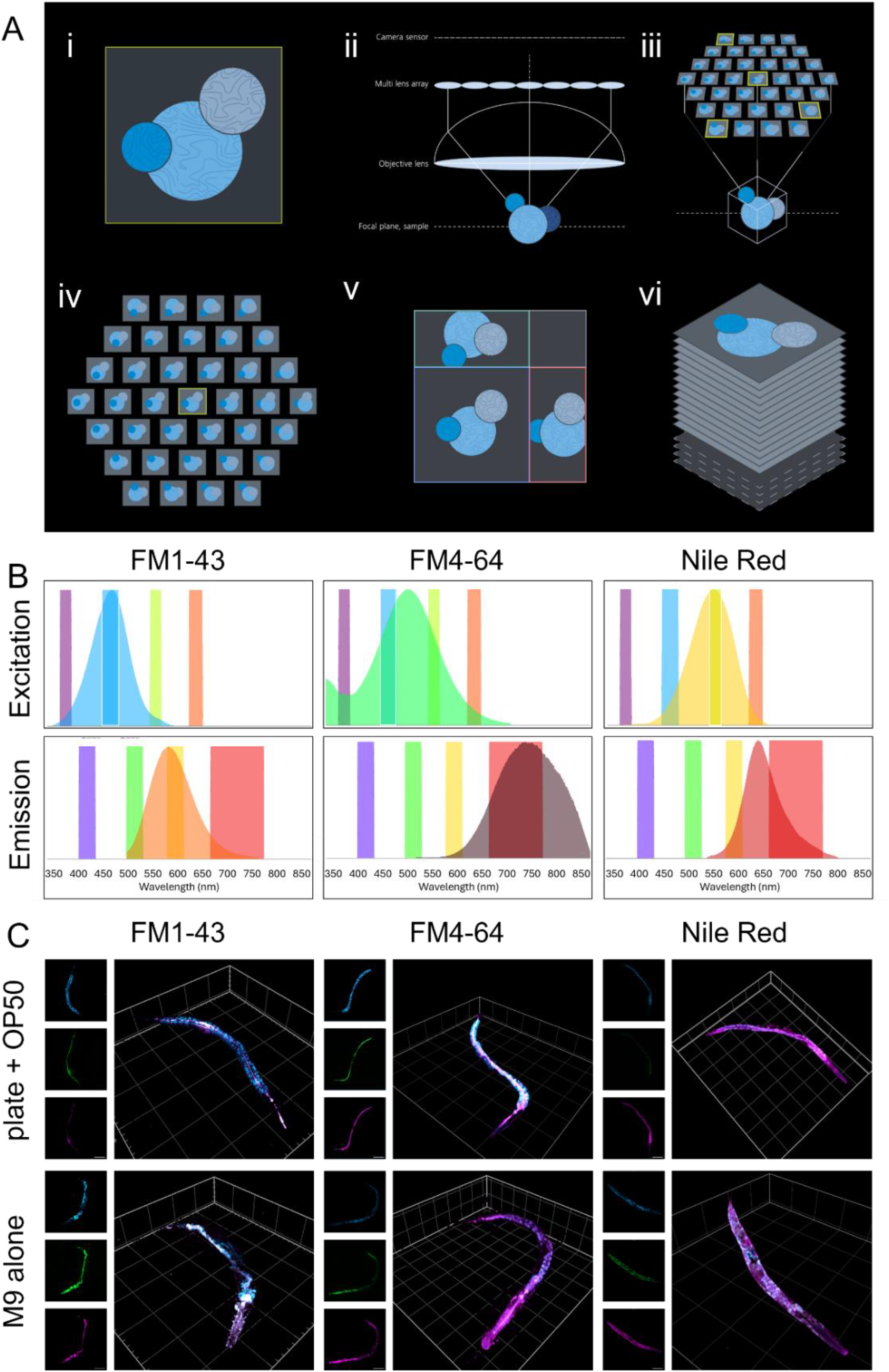
Principle of light-field volumetric imaging and dye- and delivery-dependent staining patterns in *C. elegans*. (A) Schematic of light-field image formation, adapted from Carl Zeiss Microscopy [13]. (i, ii) A specimen is imaged through a microlens array positioned between the objective and camera. (iii–v) The camera records 37 simultaneous views containing spatial and angular information, which (vi) are reconstructed by deconvolution into a three-dimensional z-stack. With the 40×/1.20 NA objective used here, the rendered stack contained 121 axial slices spanning 109 µm at 0.91 µm spacing. (B) Excitation and emission spectra of FM1-43, FM4-64 and Nile Red, plotted using FPbase [14]. (C) Representative reconstructed volumes following feeding on dye-treated E. coli OP50 or direct exposure in M9 buffer, showing dye- and delivery-dependent staining patterns. Grid spacing 100 µm (FM4-64, M9 alone, 50 µm).

### FM1-43

Distinct dye and delivery-dependent volumetric patterns were observed. FM1-43 produced combined intestinal, cuticular and vesicular labelling, with signal detected in both the green and red fluorescent channels [4]. Following feeding, the dominant signal was intestinal, consistent with uptake through ingestion of dye-labelled OP50 [9], although occasional cuticle-associated signal was also present. Following direct exposure to the FM1-43 dye, labelling was broader and included prominent vesicular structures in addition to intestinal and subtle cuticular regions. Under these conditions, FM1-43 therefore reported both internal uptake and external membrane-associated features.

### FM4-64

FM4-64 produced a related but more distinct pattern [10]. Following feeding on dyed bacteria, labelled bacterial mass was more clearly evident within the worm and intestinal signal was more readily resolved, with less cuticle staining than observed for FM1-43. By contrast, direct exposure to the FM4-64 dye produced less specific labelling dominated by vesicular and tissue-associated structures, with some lower intestinal material also visible. Under these conditions, FM4-64 therefore provided clearer visual separation between bacterial ingestion-associated signal and the broader fluorescence observed after liquid exposure.

### Nile Red

Nile Red showed a further distinct profile. Signal was distributed more broadly across nematode anatomy than with the FM dyes, including surface-associated [11] and internal regions, but remained enriched in vesicular structures [5,12]. Bacteria-fed nematodes showed relatively stronger intestinal signal, whereas direct exposure to Nile Red dye produced broader whole-worm fluorescence with less obvious intestinal enrichment. Embryos displayed reduced Nile Red signal relative to the rest of the worm. Under these conditions, Nile Red highlighted vesicular staining while retaining whole-animal context.

## Conclusions

This exploratory study was designed to compare representative volumetric staining patterns rather than quantify uptake kinetics or absolute fluorescence intensity, and no inferential statistical analysis was performed. Importantly, the acquisition is volumetric in all three spatial dimensions: with the 40× objective, 109 µm of axial range was encoded in each single-channel exposure and rendered as 121 slices at 0.91 µm spacing. The ability to capture distinct delivery-dependent whole-worm fluorescence patterns together with substantial axial depth in milliseconds highlights light-field microscopy as a rapid platform for whole-organism fluorescence mapping. Future studies can build on this workflow through targeted reporters, quantitative segmentation and time-resolved acquisition.

## Methods

### Dye delivery by feeding

For bacterial delivery, 100 µL of a 1 mM dye solution (FM1-43, FM4-64 or Nile Red; Invitrogen) was soaked onto an *E. coli* OP50 lawn and incubated overnight. Approximately 20–30 adult *C. elegans* were placed on the dyed lawn for 24 h. Nematodes were washed three times with M9 buffer and stored on ice until imaging.

### Direct dye exposure

For direct exposure, nematodes were fully suspended in the corresponding dye solution for 24 h. Nematodes were washed three times with M9 buffer and stored on ice until imaging.

### Slide preparation

Nematodes were mounted on agar pads, excess liquid removed and 5 µL of 10 mM levamisole added for immobilisation. Samples were covered with a ZEISS No.1.5 thickness coverslip.

### Light-field imaging

Live nematodes were imaged on a ZEISS Axio Observer.Z1/7 inverted microscope fitted with a ZEISS Lightfield 4D module, using a PCO camera coupled through a ZEISS Lightfield 4D adapter. Volumetric information was captured in a single exposure per channel and reconstructed using the ZEISS Lightfield 4D processing function in ZEN (∼59 s per dataset). Imaging used a C-Apochromat 40×/1.20 W Korr FCS water-immersion objective (effective NA 1.20, 1× tube lens) with a 90 HE DAPI/GFP/Cy3/Cy5 quad-band reflector (beam splitter 405/493/575/653). Three channels were acquired per experiment, where either red fluorescence emission was used for FM1-43 and Nile Red or far-red emission range was used for the FM4-64-stained samples. Fluorescence settings: (i) blue — LED 385 nm at 15%, excitation 375–395 nm, emission 410–440 nm, 50 ms; (ii) green (FM1-43) — LED 475 nm at 5%, excitation 455–483 nm, emission 499–529 nm and 579–604 nm (due to wide emission of FM1-43), 12 ms; (iii) red (Nile Red) — LED 564 nm at 20%, excitation 545–564 nm, emission 579–604 nm, 50 ms; and (iv) far-red (FM4-64) — LED 475 nm at 10%, excitation 455–483 nm, emission 659–759 nm, 50 ms. Images were recorded at 16-bit depth, 1×1 binning and 0.704 µm/pixel over a 685 µm × 1010 µm field of view (973 × 1433 pixels). With this optical configuration, each rendered dataset comprised 121 axial slices spanning 109 µm, corresponding to a reconstructed axial spacing of 0.91 µm. In a representative adult, nematode signal occupied the volume from 40.88 to 85.4 µm, an axial extent of approximately 44.5 µm. These values describe the reconstructed axial dataset generated from simultaneously acquired light-field information rather than a mechanically stepped z-stack. Reconstructed volumes are shown without further segmentation; individual worms were cropped for display only.

### Simulated confocal

Equivalent confocal acquisition times were estimated using a ZEISS Cell Discoverer 7 (Plan-Apochromat 20×/0.7) operating over a comparable cropped field of view. A single volumetric z-stack covering ∼400 × 400 µm of comparable tissue required ∼3 min. Tiling to cover the ∼700 × 1000 µm field occupied by a single nematode required a 2 × 3 raster of six such stacks, giving an estimated ∼18 min per worm. This is approximately four orders of magnitude (∼10,000-fold) slower than the summed exposure time of 112 ms across three channels used here. Estimates are provided for comparison only, matched to the field of view, channel number and axial range of the light-field acquisition, but not acquired as confocal data.

### Statistical analysis

No statistical analyses were performed.

### Reagents

#### Strains

| Strain | Genotype | Available from |
| --- | --- | --- |
| N2 | *Caenorhabditis elegans* wild type | CGC |

#### Dyes

| Reagent | Supplier | Use |
| --- | --- | --- |
| FM1-43 | Invitrogen | Membrane / vesicle labelling |
| FM4-64 | Invitrogen | Membrane / bacterial ingestion |
| Nile Red | Invitrogen | Lipid vesicular staining |

## Declaration of generative AI in manuscript

During the preparation of this work, the authors used large language models to support the organisation, language and readability of the manuscript. The authors reviewed and edited the content as needed and take full responsibility for the content of the published article.

## Data availability

Representative reconstructed volumes are shown in Figure 1. Source and reconstructed data are available from the corresponding author on reasonable request.

## Author Contributions

R.M.: conceptualisation, data curation, formal analysis, investigation, methodology, resources, visualisation, writing – review & editing. E.R.: formal analysis, methodology, resources, validation, visualisation, writing – review & editing. N.A.B.: writing – review & editing. V.T.: funding acquisition, supervision, writing – review & editing. C.H.: resources, writing – review & editing. D.R.A.W.: funding acquisition, resources, writing – review & editing. V.M.C.: conceptualisation, data curation, formal analysis, funding acquisition, investigation, methodology, project administration, resources, supervision, validation, visualisation, writing – original draft, writing – review & editing.

## Acknowledgements

We thank the University of Nottingham School of Life Sciences Imaging (SLIM) facility for imaging support and for hosting ZEISS’s demonstration of the ZEISS Lightfield 4D microscope. Some strains were provided by the CGC, which is funded by the NIH Office of Research Infrastructure Programs (P40 OD010440).

